# ModelCIF: An extension of PDBx/mmCIF data representation for computed structure models

**DOI:** 10.1101/2022.12.06.518550

**Authors:** Brinda Vallat, Gerardo Tauriello, Stefan Bienert, Juergen Haas, Benjamin M. Webb, Augustin Žídek, Wei Zheng, Ezra Peisach, Dennis W. Piehl, Ivan Anischanka, Ian Sillitoe, James Tolchard, Mihaly Varadi, David Baker, Christine Orengo, Yang Zhang, Jeffrey C. Hoch, Genji Kurisu, Ardan Patwardhan, Sameer Velankar, Stephen K. Burley, Andrej Sali, Torsten Schwede, Helen M. Berman, John D. Westbrook

## Abstract

ModelCIF (github.com/ihmwg/ModelCIF) is a data information framework developed for and by computational structural biologists to enable delivery of *Findable, Accessible, Interoperable*, and *Reusable* (*FAIR*) data to users worldwide. It is an extension of the Protein Data Bank Exchange / macromolecular Crystallographic Information Framework (PDBx/mmCIF), which is the global data standard for representing experimentally-determined, three-dimensional (3D) structures of macromolecules and associated metadata. ModelCIF provides an extensible data representation for deposition, archiving, and public dissemination of predicted 3D models of proteins. The PDBx/mmCIF framework and its extensions (*e.g*., ModelCIF) are managed by the Worldwide Protein Data Bank partnership (wwPDB, wwpdb.org) in collaboration with relevant community stakeholders such as the wwPDB ModelCIF Working Group (wwpdb.org/task/modelcif). This semantically rich and extensible data framework for representing computed structure models (CSMs) accelerates the pace of scientific discovery. Herein, we describe the architecture, contents, and governance of ModelCIF, and tools and processes for maintaining and extending the data standard. Community tools and software libraries that support ModelCIF are also described.

## INTRODUCTION

### Brief History of Computed Structure Models (CSMs)

Protein Data Bank (PDB) is the single global repository for three-dimensional (3D) structures of biological macromolecules determined experimentally using macromolecular crystallography (MX), nuclear magnetic resonance (NMR) spectroscopy, and electron microscopy (3DEM). It was established in 1971 as the first open-access digital data resource in biology with seven protein structures [1, 2]. At the time of writing, the archive contained ~200,000 structures of proteins, nucleic acids and their complexes with one another and with small-molecule ligands (*e.g*., approved drugs, investigational agents, enzyme cofactors). This metric is a testament to the collective efforts and technological advances made by structural biologists working on all inhabited continents. It also highlights a daunting reality—that 99% of protein structure space remains unexplored by experimental methods. Inspired by the work of Anfinsen [3], computational structural biologists began trying to predict the 3D structure of a protein from its amino sequence.

Two distinct approaches for protein structure prediction [4] have been pursued (Figure 1). The first approach is template-based structure prediction (also known as homology modeling or comparative modeling), in which the structure of an unknown protein (target) is modeled computationally based on the similarity of its amino acid sequence to that of a protein with a known structure (template). Homology modeling is generally successful when template structures from the PDB can be identified and accurately aligned to the target sequence. The second approach is template-free structure prediction, also known as *ab initio* or *de novo* modeling, which can be applied when reliable structural templates are not available for the protein of interest. In recent years, intramolecular residue-residue contact predictions based on coevolution data [5] have been successfully applied for template-free structure prediction [6].

**Figure 1.**
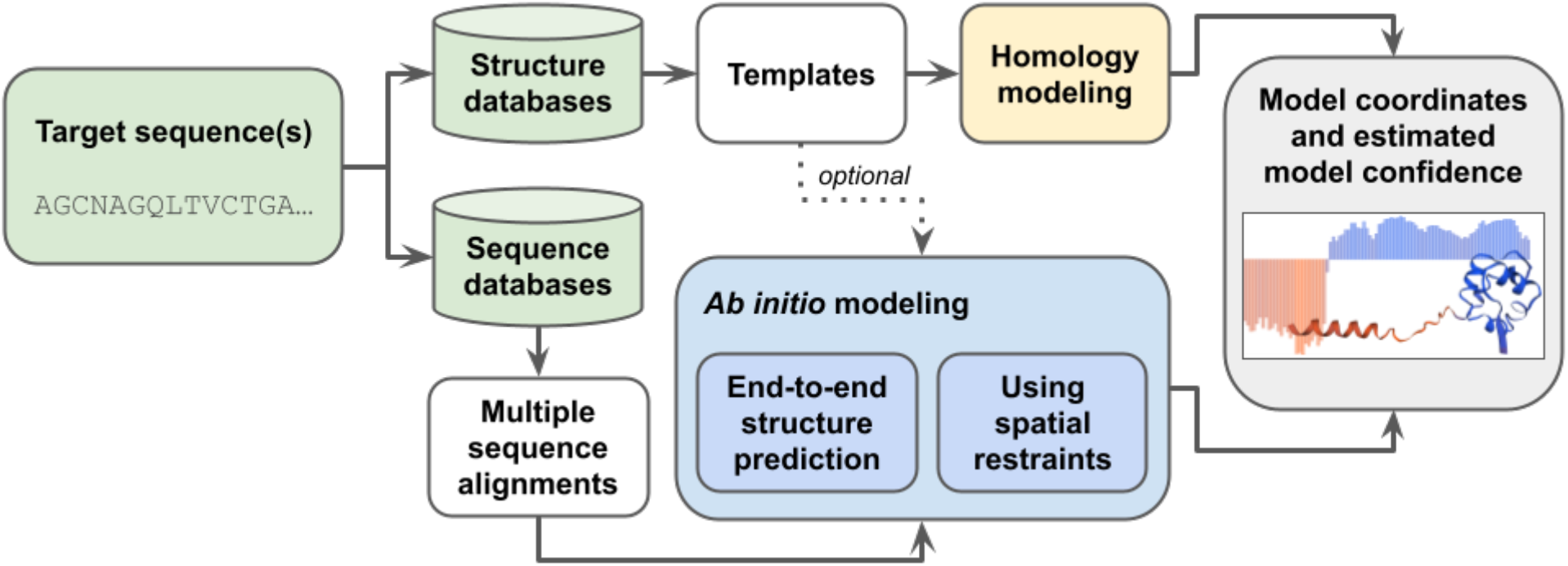
Schematic representation of modeling methods using target sequence(s), structure databases (*e.g*., PDB), and sequence databases (*e.g*., Uniclust30 [45]) as input to produce CSMs and estimates of prediction confidence. Homology modeling uses specific templates as its main input, while *ab initio* methods work without templates. Commonly used *ab initio* methods rely on multiple sequence alignments, which are either used directly as input for end-to-end structure prediction or processed to extract spatial restraints used to generate CSMs.

Several automated software tools and web servers support template-based or template-free structure prediction, including, but not limited to, SWISS-MODEL [7], Modeller [8], ROSETTA [9], I-TASSER [10], QUARK [11], AlphaFold2 [12], and RoseTTAFold [13, 14]. In the Critical Assessment of Techniques for Protein Structure Prediction (CASP14) challenge conducted in 2020 [15], AlphaFold2 demonstrated unprecedented levels of success, an achievement largely enabled by breakthroughs applying machine learning (ML) approaches to protein structure prediction. Following CASP14, RoseTTAFold, another ML-based method, was developed by David Baker and colleagues at the University of Washington, which was recently applied in combination with AlphaFold2 to predict the structures of hetero-dimeric complexes of eukaryotic proteins [14]. These ML-based structure prediction methods have proven highly successful and are now capable of generating computed structure models (CSMs) with accuracies comparable to that of lower-resolution experimentally-determined structures [16].

Paralleling advances in protein structure prediction methodologies, data resources were established to provide open access to modeled structures. SWISS-MODEL Repository [17] and ModBase [18] house millions of CSMs of proteins generated using SWISS-MODEL or Modeller, respectively. In addition, the ModelArchive, developed at the Swiss Institute of Bioinformatics (SIB, https://www.modelarchive.org/), was created to archive and provide stable digital object identifiers (DOIs) for CSMs referenced in publications. ModelArchive includes CSMs which were stored in the PDB before 2006 and has been accepting new depositions since 2013. At the time of writing, the AlphaFold Protein Structure Database (AlphaFold DB) [19] held more than 200 million protein CSMs generated by AlphaFold2. They are freely available and represent virtually all of the protein sequences cataloged in UniProtKB [20].

### Significance of data standards in archiving scientific data

Data standards are technical specifications describing the semantics, logical organization, and physical encoding of data and associated metadata. They serve as the foundation for collecting, processing, archiving, and distributing data in a standard format and promoting the *FAIR* (*Findable, Accessible, Interoperable and Reusable*) principles emblematic of responsible data management in the modern era [21]. In addition to representing the results of a scientific investigation, additional metadata (such as software, authors, citations, references to external data) may be required to support data exchange among different stakeholders, including data generators, archives, and data consumers. If a consistent mechanism is utilized to store such information, it can be shared using common software, agnostic of the data provider, enabling better interoperation among resources and facilitating data search, retrieval, and reuse. Involving community experts in developing and subsequently extending data standards ensures that they are readily adopted by the community and facilitates continuous update of the standards as the field evolves.

### History of PDBx/mmCIF data standard for representing macromolecular structures

One of the earliest archival formats in structural biology is the legacy PDB format [22]. Developed in the 1970s, it is human and machine readable, easy to parse, and remained the PDB standard exchange format for over forty years. However, it has several drawbacks, including fixed field widths, column positions, and metadata format, which posed severe limitations for archiving large macromolecular structures, data validation, and future expansion to support newer experimental methods.

In 1990, the Crystallographic Information Framework (CIF) [23] was adopted by the International Union of Crystallography (IUCr) as a community data standard to describe small-molecule X-ray diffraction studies. Later, in 1997, the IUCr approved the mmCIF data standard [24] to support MX experiments. The original mmCIF data standard was subsequently extended by the PDB to support other experimental methods (*e.g*., NMR, 3DEM), and to create the PDBx/mmCIF data dictionary [25, 26]. In 2014, this standard was adopted by the worldwide PDB (wwPDB, wwpdb.org) [2, 27] as the master format for the PDB archive. The framework describing PDBx/mmCIF is regulated by Dictionary Definition Language 2 (DDL2), a generic language that supports construction of dictionaries composed of data items grouped into categories [28]. DDL2 supports primary data types (*e.g*., integers, real numbers, text), boundary conditions, controlled vocabularies, and linking of data items together to express relationships (*e.g*., parent–child relationships). Additionally, software tools have been developed to manage the PDBx/mmCIF dictionary (mmcif.wwpdb.org/docs/software-resources.html). PDBx/mmCIF overcame the limitations of the legacy PDB format and has been extended to represent small-angle solution scattering data [29] and integrative structure models [30].

### History of ModelCIF and the wwPDB ModelCIF Working Group

Initial efforts to extend PDBx/mmCIF to support CSMs began in 2001 with creation of the MDB dictionary [31]. In 2006, the outcomes of a Workshop organized by the Research Collaboratory for Structural Bioinformatics (RCSB) PDB at Rutgers included recommendations to build a common portal for accessing structural models and develop data standards to support CSMs [32]. The Protein Model Portal (PMP) [33] was created at SIB in collaboration with the Protein Structure Initiative (PSI) Structural Biology Knowledgebase [34]. A collaborative project between RCSB PDB and SIB was initiated in 2016 to create data standards that represent CSMs in the PMP and the ModelArchive. These data standards were designed as an extension of PDBx/mmCIF to facilitate interoperation with PDB data. The first set of ModelCIF definitions was released on GitHub in 2018 (github.com/ihmwg/ModelCIF).

The ModelCIF Working Group (WG) was established in 2021 as a collaboration between the wwPDB partners (RCSB PDB, Protein Data Bank in Europe (PDBe), Protein Data Bank Japan (PDBj), Electron Microscopy Data Bank (EMDB), and Biological Magnetic Resonance Bank (BMRB)) and domain experts in computational structural biology (wwpdb.org/task/modelcif). In addition to wwPDB members, the WG includes representatives from ModelArchive, SWISS-MODEL, Genome3D [35], ModBase, I-TASSER, AlphaFold database, AlphaFold2/DeepMind, and RoseTTAFold. The WG is involved in development and maintenance of the ModelCIF data standard for representing and archiving CSMs and promotes its adoption across the computational biology community. The WG also promotes development of software tools supporting ModelCIF, such as the python-modelcif software library (github.com/ihmwg/python-modelcif). Feedback to the WG *via* email is welcome.

## RESULTS AND DISCUSSION

### Data definitions reused from PDBx/mmCIF

ModelCIF is an extension of PDBx/mmCIF for describing the specific set of attributes and metadata associated with macromolecular structures modeled by solely computational methods. In developing ModelCIF, various core PDBx/mmCIF dictionary definitions have been reused. These include representation of small-molecules, polymeric macromolecules, biomolecular complexes, and their atomic coordinates, as well as related metadata definitions about modeling software used, bibliographic citations, and author names (Figure 2).

**Figure 2.**
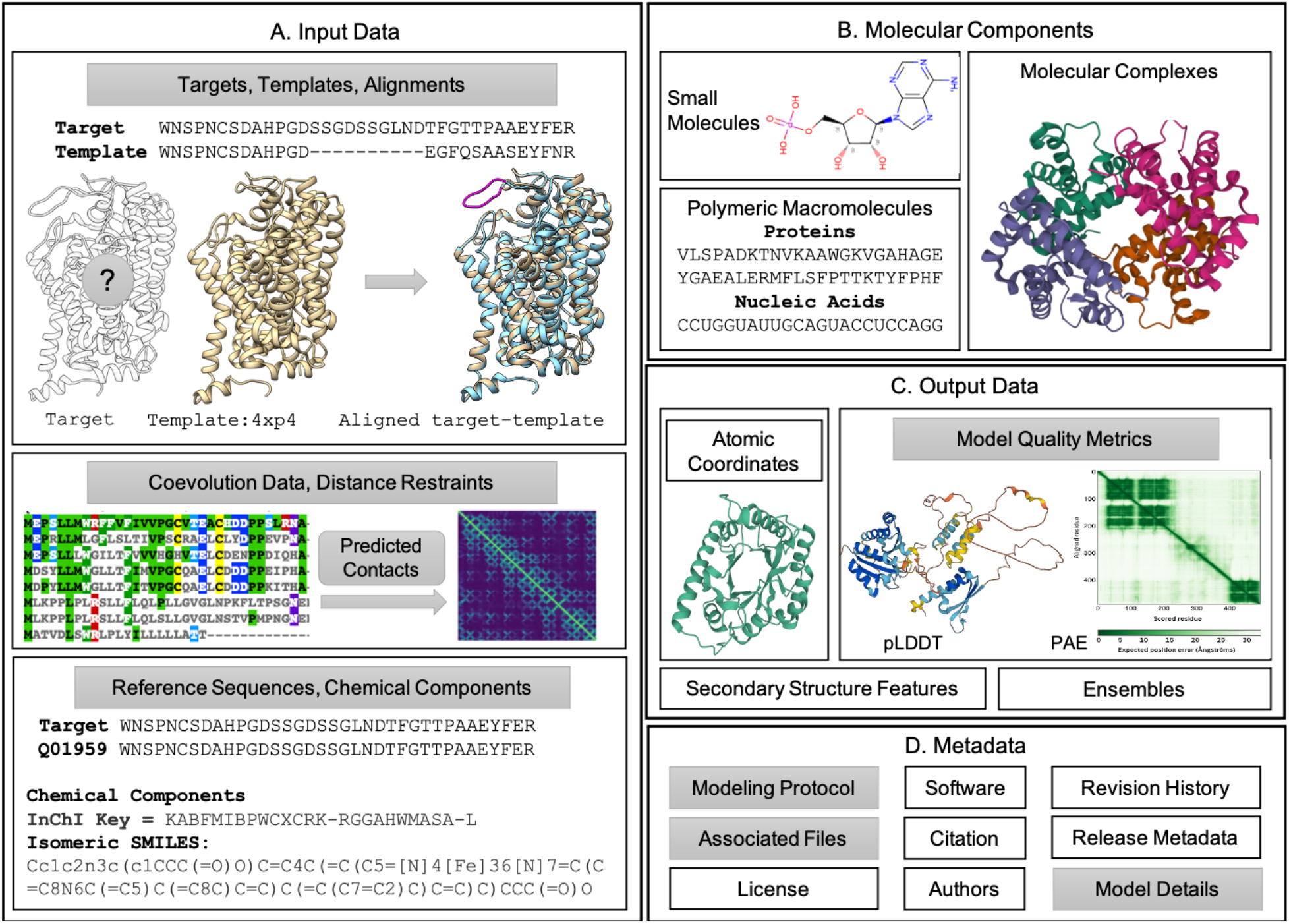
Schematic representation of the data specifications in ModelCIF. Definitions reused from PDBx/mmCIF are shown in white boxes (*e.g*., Atomic Coordinates) and the newly added definitions are shown in gray boxes (*e.g*., Model Quality Metrics). (A) Descriptions are provided for input data used in template-based and template-free modeling. (B) Representations of molecular components are retained from PDBx/mmCIF. (C) Definitions for atomic coordinates, secondary structure features, and ensembles are taken from PDBx/mmCIF; descriptions of local and global CSM quality metrics are defined in ModelCIF. (D) Several metadata definitions from PDBx/mmCIF are reused. New metadata definitions regarding modeling protocol, CSM classification (*ab initio*, homology, *etc*.) and descriptions of associated files are included in ModelCIF.

### ModelCIF data definitions

Given the variety of existing modeling methods, ModelCIF aims to be flexible regarding data representation. To fulfill this goal, new data categories were introduced to: (i) store input and intermediate results that are of relevance for existing methods; (ii) provide estimates of local and global CSM confidence; (iii) describe steps used to generate CSMs; and (iv) refer to data stored in associated files. New ModelCIF definitions are summarized in Figure 2.

In addition to CSM atomic coordinates, two sets of data items are mandatory: (i) details regarding modeled targets and (ii) list of CSMs included in the file. New definitions are provided for capturing information pertaining to the origin of modeled molecular entities. This feature is particularly useful for cross-referencing to external databases for macromolecular sequences (*e.g*., UniProtKB) and small-molecules (*e.g*., PubChem [36], ChEBI[37]). Definitions to include small-molecules that are not already specified in the wwPDB chemical component dictionary (CCD) [38] are also provided.

In ModelCIF, CSMs can be combined into groups that may belong to an ensemble (or cluster). Structural assemblies must be homogeneous (*i.e*., every CSM in an entry must have identical composition and structural elements). Each CSM can be classified as “homology model”, *“ab initio* model” or “other” if neither descriptor is appropriate. The “homology model” category is used for any modeling method (including comparative modeling and protein threading) where the main inputs for generating the CSM are sequence alignments to templates. CSMs generated without templates (or where templates are not considered dominant inputs) are classified as “*ab initio* model” (including fragment sampling and ML-based methods).

Homology modeling methods, as used by SWISS-MODEL and Modeller for example, typically consist of three steps: (i) template identification; (ii) target-template alignment; and (iii) atomic coordinate generation. ModelCIF includes data categories to store the most relevant intermediate results in a standardized way, including a summarized version of the template search results with cross-references to relevant structure databases (*e.g*., PDB) and detailed information regarding template structures and target-template alignments used for modeling.

*Ab initio* methods start from sequence information without relying on structural templates. Methods such as I-TASSER generate CSMs using folding simulations guided by deep learning predicted spatial restraints extracted from multiple sequence alignments (MSAs) and corresponding co-evolutionary features. The spatial restraints from deep learning predictors could be residue-residue contacts, distances, dihedral angles, torsion angles, or hydrogen-bonding networks. ModelCIF enables storage of MSAs, homologous templates (optionally used as input structures for *ab initio* methods), and derived spatial restraints, used by *ab initio* folding simulations to model CSMs. ML-based *ab initio* methods such as AlphaFold2 and RoseTTAFold do not rely on features extracted from templates or MSAs, but can instead use them as raw input to an “end-to-end” neural network that directly generates the atomic coordinates. Consequently, ModelCIF allows for inclusion of simplified descriptions of relevant input data and intermediate results. ModelCIF can also store information about sequence databases used to construct MSAs (including versions and download URLs) and minimal details regarding any input structures utilized.

While CSMs generated with the newest techniques have become increasingly accurate, it is critical that they are accompanied by estimates of model quality (or prediction confidence). Quality estimates are used to evaluate models and assess their suitability for specific downstream applications. ModelCIF includes flexible support to define any number of quality assessment values. These are classified according to how they are to be interpreted (*e.g*., probabilities, distances, energies) or as a prediction of the similarity to the correct structure according to well defined metrics such as the TM-score [39] or lDDT [40]. Quality estimate values can be provided globally per CSM and locally per residue, to identify high- and low-quality regions, and per residue-pair, to enable assessment of contacts and domain orientations.

To facilitate reproducibility of structure prediction and to acknowledge use of publicly available software and web services, ModelCIF allows inclusion of generic definitions describing modeling protocols. Minimally, such definitions may include a free text description of the modeling protocol as a single step. Ideally, however, they connect input data and software, including software parameters, through intermediate results, to the final CSM. To keep data file sizes manageable, ModelCIF provides metadata definitions supporting description of one or more associated files. The data content of associated files can be large intermediate results, such as MSAs or quality estimates for residue-pairs. A variety of generic file formats are allowed for associated files.

### Supporting software tools and resources

Table 1 provides a list of software tools and CSM resources that support ModelCIF. Additional details concerning these tools and resources are included in the supplementary material.

**Table 1.**
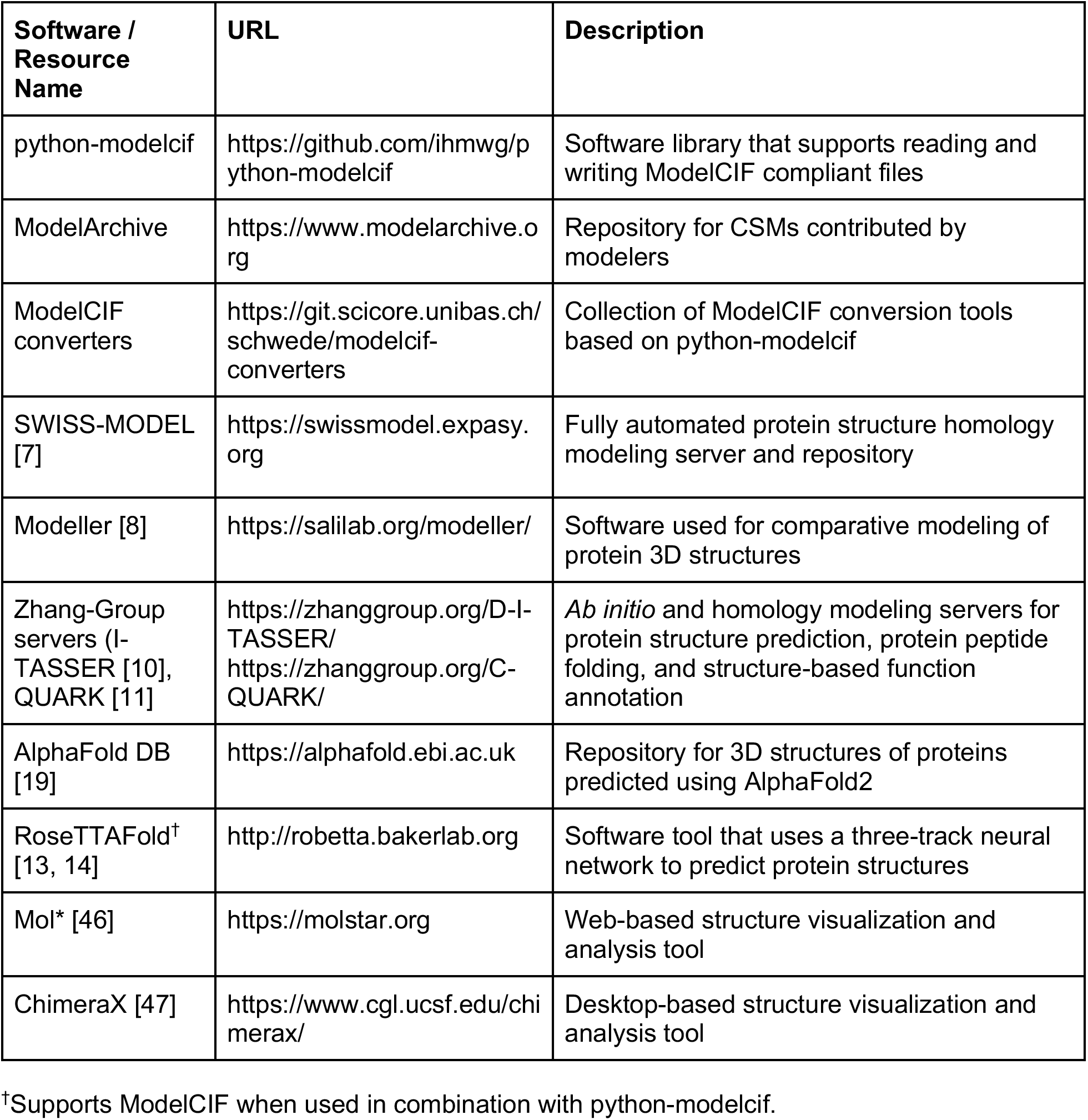
Software tools and CSM resources supporting ModelCIF.

### Advantages of ModelCIF

The value and benefits of ModelCIF are most readily recognized through its support for the *FAIR* principles. ModelCIF provides foundational data standards for archiving CSMs, making them freely available, and enabling seamless data exchange. Moreover, extending PDBx/mmCIF to establish ModelCIF as a data standard in its own right provides the following advantages: (a) existing definitions in PDBx/mmCIF for representing the atomic structures of biological macromolecules, small-molecules, and molecular complexes can be reused; (b) software tools developed to support PDBx/mmCIF can be reused and extended to support the extension; and (c) the extension facilitates interoperation with other structural biology data resources (*e.g*., PDB). For example, recent updates to the RCSB.org web portal to include >1,000,000 CSMs available freely from AlphaFoldDB and the ModelArchive was facilitated by ModelCIF [41]).

## CONCLUSION AND PERSPECTIVES

Computational structural biology is rapidly advancing before our eyes as a discipline. During manuscript preparation, Meta AI announced the development of their own ML-based method for protein structure prediction and used it to generate more than 600 million CSMs that are now publicly available [42]. It is also likely that additional open-access resources distributing CSMs of proteins will emerge before this paper appears in print. Ideally, every one of these newly established databases of predicted structures will embrace the ModelCIF data standard for deposition, archiving, and dissemination of CSMs. The wwPDB ModelCIF Working Group is committed to maintaining and updating the data standard as new approaches to computational structure modeling of biological macromolecules emerge and are validated. The wwPDB is also supporting community efforts, such as the 3D-Beacons network [43], to encourage adoption of common data standards and facilitate access to 3D-structure information.

Looking ahead, CSMs of large, intricately folded ribonucleic acid (RNA) chains may be of particular importance to basic and applied researchers working across fundamental biology, biomedicine, biotechnology/bioengineering, and the energy sciences. Progress in this field was reported in 2021 by researchers at Stanford University using their Atomic Rotationally Equivariant Scorer (ARES) system [44]. As for protein structure prediction, community-organized blind challenges will be important in accelerating technical developments in this area. CASP15 (predictioncenter.org/casp15/), underway at the time of writing, includes structure prediction of RNA molecules.

## ACKNOWLEDGEMENTS

The authors thank the tens of thousands of researchers worldwide who enable computational structure modeling of proteins by depositing ~200,000 experimentally-determined structures to the PDB since 1971. We also gratefully acknowledge contributions to the PDBx/mmCIF data standard made by past members of Worldwide Protein Data Bank partner organizations (RCSB PDB, PDBe, PDBj, EMDB, and BMRB) and members of the structural biology community. G.T., S.B., and T.S. acknowledge the contributions of Andrew Mark Waterhouse and Dario Behringer to ModelArchive and support of ModelCIF.

