## Supplementary text for "ModelCIF: An extension of PDBx/mmCIF data representation for computed structure models"

### SUPPLEMENTARY MATERIAL

#### *Methods*

##### *Extending PDBx/mmCIF for computed structure models*

ModelCIF is an extension of PDBx/mmCIF [1, 2] for describing macromolecular structures modeled by computational methods. The definitions in ModelCIF are the result of collaborations and discussions among the ModelCIF working group members (wwpdb.org/task/modelcif) and are based on examples of computed structure models (CSMs) obtained from modeling resources such as SWISS-MODEL repository [3], RoseTTAFold [4, 5], ModelArchive (https://www.modelarchive.org), ModBase [6], I-TASSER [7, 8], and AlphaFoldDB [9]. These collaborative efforts have culminated in the creation of version 1.4.3 of the ModelCIF dictionary extension released in September 2022 (https://mmcif.wwpdb.org/dictionaries/mmcif_ma.dic/Index/). ModelCIF is licensed under Creative Commons Zero v1.0 Universal (CC0 1.0) license.

##### *ModelCIF dictionary development (GitHub)*

The ModelCIF dictionary is developed as a collaborative project on a public GitHub repository (https://github.com/ihmwg/ModelCIF). The complete ModelCIF dictionary extension file is available under the distribution (i.e., "dist") directory (mmcif_ma.dic) within the repository. This complete and distributable form of the extension is constructed from the concatenation of the individual source dictionary components found under the "base" directory (https://github.com/ihmwg/ModelCIF/tree/master/base); thus, these "base" files are what are directly modified by maintainers, which are then combined to generate the complete distributable extension file (using the "build" scripts in https://github.com/ihmwg/ModelCIF/tree/master/scripts). Any additions or modifications arising from the discussions in the ModelCIF WG are incorporated in the source dictionary files on GitHub, followed by a full update and release of the dictionary. Whenever ModelCIF is updated, the latest PDBx/mmCIF dictionary is pulled from the corresponding Git repository (https://github.com/wwpdb-dictionaries/mmcif_pdbx) to ensure compatibility and compliance with the latest PDBx/mmCIF definitions. Importantly, upon each update, the ModelCIF version number is incremented and metadata regarding the revision history is included.

##### *ModelCIF validation and maintenance (GitHub, mmCIF resources website, PDBx/mmCIF dictionary tools)*

Following the introduction of any new modifications and version updates to ModelCIF, the resulting fully-formed dictionary undergoes a series of quality checks to ensure the dictionary is formatted properly, is free of any syntax errors, and successfully validates a set of example model files. Some of these checks are performed while building the concatenated dictionary file from the source "base" files described above. In addition, PDBx/mmCIF software tools (e.g., https://github.com/rcsb/cpp-dict-pack) are used to further validate the updated dictionary. Only upon successful passing of these validation tests is the new version of the dictionary extension released on GitHub and published on the PDBx/mmCIF resources website, mmcif.wwpdb.org (i.e., https://mmcif.wwpdb.org/dictionaries/mmcif_ma.dic/Index/).

### ***Results and discussion***

#### *Supporting software tools and resources*

The following software tools and CSM resources support ModelCIF (also listed in Table 1 in the main text).

**python-modelcif**: The python-modelcif library (https://github.com/ihmwg/python-modelcif) provides a mechanism to describe a structural model as a hierarchy of Python objects. These objects capture the same information as in the ModelCIF dictionary, and can be written out to or read in from ModelCIF files in either mmCIF or BinaryCIF [10] format. The library automatically handles the various object identifiers used in the ModelCIF dictionary, mapping them naturally to interrelationships between Python objects, and mapping controlled vocabulary terms defined in the dictionary to Python subclasses. The library thus generates conformant ModelCIF files "by construction", although it is also able to read in and validate files against the ModelCIF dictionary. Convenience Python application programming interfaces (APIs) are also provided to add additional metadata to output files, such as citation information from PubMed, reference sequence information from UniProtKB [11], and metadata about associated files. The library is designed such that it can be embedded in a modeling or visualization package to support input or output of ModelCIF files, or it can be used standalone, and it is available under the permissive MIT open-source license.

**ModelArchive:** ModelArchive supports the submission of ModelCIF files. The information necessary for deposition is extracted from the metadata in the ModelCIF file. ModelCIF files are also mandatory for the deposition of large datasets. Current datasets include structural predictions of core eukaryotic protein complexes generated by RoseTTAFold [4, 5] in combination with AlphaFold2 [12] (ma-bak-cepc) [5], structural and functional predictions using ColabFold in combination with FoldSeek (ma-coffe-slac), structural predictions for *Mycobacterium tuberculosis* proteins of relevance for antibiotic resistance obtained using a combination of SWISS-MODEL [13], AlphaFold-Multimer and small molecule docking (ma-tbvar3d), and AlphaFold2 predictions of *Sphagnum divinum* (ma-ornl-sphdiv) [14] and the African swine fever virus (ma-asfv-asfvg). To assist such depositions and generate correctly formatted ModelCIF files, the ModelArchive team developed tools (https://git.scicore.unibas.ch/schwede/modelcif-converters) based on python-modelcif and OpenStructure [15]. ModelArchive furthermore takes advantage of cross-references to UniProtKB [11] included in ModelCIF files to connect these models within the 3D-Beacons network [16], which provides unified data access to models from various model providers.

**SWISS-MODEL:** The ModelCIF format is well suited to provide a single output file for homology models generated by SWISS-MODEL [13]. Experimental support for ModelCIF files is available for interactive SWISS-MODEL projects and includes information on templates and alignments used as well as model quality assessments. Work is in progress to add ModelCIF support to the SWISS-MODEL Repository [3].

**Modeller**: The Modeller [17] comparative modeling pipeline supports output of models in ModelCIF compliant format, including information on the template(s) and alignment(s) used.

**Zhang-Group servers**: Most protein structure prediction servers from Zhang-Group now support ModelCIF. For example, I-TASSER related servers [7, 8], QUARK related servers [18, 19], and the LOMETS [20] server can provide the final predicted model in both PDB format [21] and ModelCIF format. In addition to the model coordinates information, the ModelCIF-formatted output also includes multiple sequence alignments, structural templates, deep learning predicted spatial restraints, and both global and per-residue model quality assessment scores.

**AlphaFoldDB:** AlphaFoldDB [9] is a database of protein structures predicted using AlphaFold2 [12] and as of October 2022 it contains predicted monomer structures for over 214 million UniProtKB [11] sequences. AlphaFoldDB uses ModelCIF as the primary data format. It is used for visualization of the predicted structures on the website and distribution of the data to users. Since the July 2022 release, all AlphaFoldDB entries are ModelCIF compliant. AlphaFoldDB ModelCIF files provide the 3D coordinate data together with the predicted LDDT (pLDDT) scores [22], secondary structure information, a list of template structures provided as extra input to the AlphaFold2 model during structure prediction, and metadata about the sequence extracted from UniProtKB. The Predicted Aligned Error (PAE) is currently provided in a separate JavaScript Object Notation (JSON) file. A subset of the data available in the ModelCIF file is also provided in the PDB format for every AlphaFoldDB entry, but only for compatibility with legacy tools. New data and metadata will be added only in the ModelCIF files going forward. AlphaFoldDB models are available via the 3D-Beacons network [16], which provides unified data access to models from various model providers.

**RoseTTAFold:** RoseTTAFold [4, 5] is a three-track neural network model which enables rapid prediction of atomic structures for single protein chains as well as protein-protein complexes. RoseTTAFold is the default modeling protocol in the Robetta protein structure prediction server (http://robetta.bakerlab.org/) [23], but it can also be deployed locally (https://github.com/RosettaCommons/RoseTTAFold). In both cases, the user gets the results in the PDB format which can then be converted into ModelCIF using the python-modelcif library.

**Structure visualization tools:** CSMs can be visualized using structure visualization tools such as Mol* [24] and ChimeraX [25]. Both Mol* and ChimeraX support ModelCIF. In addition to visualizing atomic structures, local model quality score-based coloring is supported. ChimeraX also supports visualization of target-template alignments described in ModelCIF.

***References***

[1] Westbrook JD, Young JY, Shao C, Feng Z, Guranovic V, Lawson C, et al. PDBx/mmCIF Ecosystem: Foundational semantic tools for structural biology. J Mol Biol. 2022;434:167599.

[2] Westbrook J, Henrick K, Ulrich EL, Berman HM. 3.6.2 The Protein Data Bank exchange data dictionary. In: Hall SR, McMahon B, editors. International Tables for Crystallography. Dordrecht, The Netherlands: Springer; 2005. p. 195-8.
[3] Bienert S, Waterhouse A, de Beer TA, Tauriello G, Studer G, Bordoli L, et al. The SWISS-MODEL Repository-new features and functionality. Nucleic Acids Res. 2017;45:D313-D9.
[4] Baek M, DiMaio F, Anishchenko I, Dauparas J, Ovchinnikov S, Lee GR, et al. Accurate prediction of protein structures and interactions using a three-track neural network. Science. 2021;373:871-6.
[5] Humphreys IR, Pei J, Baek M, Krishnakumar A, Anishchenko I, Ovchinnikov S, et al. Computed structures of core eukaryotic protein complexes. Science. 2021;374:eabm4805.
[6] Pieper U, Webb BM, Dong GQ, Schneidman-Duhovny D, Fan H, Kim SJ, et al. ModBase, a database of annotated comparative protein structure models and associated resources. Nucleic Acids Res. 2014;42:D336-46.
[7] Yang J, Yan R, Roy A, Xu D, Poisson J, Zhang Y. The I-TASSER Suite: protein structure and function prediction. Nat Methods. 2015;12:7-8.
[8] Zheng W, Zhang C, Li Y, Pearce R, Bell EW, Zhang Y. Folding non-homologous proteins by coupling deep-learning contact maps with I-TASSER assembly simulations. Cell Rep Methods. 2021;1.
[9] Varadi M, Anyango S, Deshpande M, Nair S, Natassia C, Yordanova G, et al. AlphaFold Protein Structure Database: massively expanding the structural coverage of protein-sequence space with high-accuracy models. Nucleic Acids Res. 2022;50:D439-D44.
[10] Sehnal D, Bittrich S, Velankar S, Koca J, Svobodova R, Burley SK, et al. BinaryCIF and CIFTools-Lightweight, efficient and extensible macromolecular data management. PLoS Comput Biol. 2020;16:e1008247.
[11] UniProt Consortium. UniProt: the universal protein knowledgebase in 2021. Nucleic Acids Research. 2021;49:D480-D9.
[12] Jumper J, Evans R, Pritzel A, Green T, Figurnov M, Ronneberger O, et al. Highly accurate protein structure prediction with AlphaFold. Nature. 2021;596:583-9.
[13] Waterhouse A, Bertoni M, Bienert S, Studer G, Tauriello G, Gumienny R, et al. SWISS-MODEL: homology modelling of protein structures and complexes. Nucleic Acids Res. 2018;46:W296-W303.
[14] Gao M, Coletti M, Davisdon RB, Prout R, Abraham S, Hernandez B, et al. Proteome-scale Deployment of Protein Structure Prediction Workflows on the Summit Supercomputer. IEEE International Parallel and Distributed Processing Symposium Workshops (IPDPSW), 20222022. p. 206-15.
[15] Biasini M, Schmidt T, Bienert S, Mariani V, Studer G, Haas J, et al. OpenStructure: an integrated software framework for computational structural biology. Acta Crystallographica Series D. 2013;69:701-9.
[16] Varadi M, Nair S, Sillitoe I, Tauriello G, Anyango S, Bienert S, et al. 3D-Beacons: Decreasing the gap between protein sequences and structures through a federated network of protein structure data resources. GigaScience. in press.
[17] Sali A, Blundell TL. Comparative protein modelling by satisfaction of spatial restraints. J Mol Biol. 1993;234:779-815.
[18] Mortuza SM, Zheng W, Zhang C, Li Y, Pearce R, Zhang Y. Improving fragment-based ab initio protein structure assembly using low-accuracy contact-map predictions. Nature communications. 2021;12:5011.
[19] Zheng W, Li Y, Zhang C, Zhou X, Pearce R, Bell EW, et al. Protein structure prediction using deep learning distance and hydrogen-bonding restraints in CASP14. Proteins : Structure, Function, and Genetics. 2021;89:1734-51.
